# Inpatient mobility to predict hospital-onset *Clostridium difficile*: a network approach

**DOI:** 10.1101/404160

**Authors:** Kristen Bush, Hugo Barbosa, Samir Farooq, Samuel J. Weisenthal, Melissa Trayhan, Robert J. White, Gourab Ghoshal, Martin S. Zand

## Abstract

With hospital-onset *Clostridium difficile* Infection (CDI) still a common occurrence in the U.S., this paper examines the relationship between unit-wide CDI susceptibility and inpatient mobility and creates a predictive measure of CDI called “Contagion Centrality”. A mobility network was constructed using two years of patient electronic health record (EHR) data within a 739-bed hospital (Jan. 2013 - Dec. 2014; n=72,636 admissions). Network centrality measures were calculated for each hospital unit (node) providing clinical context for each in terms of patient transfers between units (edges). Daily unit-wide CDI susceptibility scores were calculated using logistic regression and compared to network centrality measures to determine the relationship between unit CDI susceptibility and patient mobility. Closeness centrality was a statistically significant measure associated with unit susceptibility (p-value < 0.05), highlighting the importance of incoming patient mobility in CDI prevention at the unit-level. Contagion Centrality (CC) was calculated using incoming inpatient transfer rates, unit-wide susceptibility of CDI, and current hospital CDI infections. This measure is statistically significant (p-value <0.05) with our outcome of hospital-onset CDI cases, and captures the additional opportunities for transmission associated with inpatient transfers. We have used this analysis to create an easily interpretable and informative clinical tool showing this relationship and risk of hospital-onset CDI in real-time. Quantifying and visualizing the combination of inpatient transfers, unit-wide risk, and current infections help identify hospital units at risk of developing a CDI outbreak, and thus provide clinicians and infection prevention staff with advanced warning and specific location data to concentrate prevention efforts.

## 1. Introduction

The U.S. sees nearly 500,000 cases of hospital-onset *Clostridium difficile* infection (CDI) each year, resulting in approximately 29,000 deaths [1]. Infection prevention teams aim to reduce patient contact with CDI spores in the hospital environment through bleach-based or terminal cleaning of rooms, contact precautions and PPE usage by clinicians and healthcare workers, increased hand hygiene compliance, and antibiotic stewardship programs [2]. Despite these efforts, CDI is still a prominent healthcare-associated infection (HAI) in many facilities, suggesting additional factors may be facilitating transmission.

Prevention efforts to reduce exposure in the patients’ immediate environment are well established[2, 3], yet little has been done to analyze the movement, or mobility, of patients between units. Inpatient mobility (transfers) can increase contact opportunities, and make risk of CDI transmission more difficult to measure. Current surveillance methods such as contact tracing begin to scratch the surface when examining contact opportunities [4], but are considered to be retrospective and do not necessarily prevent future outbreaks stemming from other sources of infection[5]. To build upon such a method and give it a prospective aspect, it is essential to examine indirect contact opportunities that may be posed from patient mobility, as well - i.e. items touched by providers or patients such as linens, beds, equipment, and other surfaces in units. The most simplistic way to do this without attempting to track items patients and providers contact is to assume broad environmental exposure based on physical location (inpatient units). In hospitals, patients are grouped into inpatient units and transferred *x* number of times within their admission, and each inpatient transfer allows for additional opportunities for exposure and contamination of a once-clean environment.

To examine patient mobility, we use graph theory, the mathematical analysis of networks that allows us to construct a network of movement how hospital units are connected by patient movement between them[6]. Network analysis has been previously used to examine intra-hospital transfers and ambulatory care[7, 8, 9], but limited studies of inter-hospital mobility[10, 11]. Grouping patients by their location in a hospital gives us the opportunity to examine susceptibility and risk of CDI from a population perspective, and calculation of network centrality[6] provides us with context of how inpatient units are connected in our hospital via patient transfers. Individual patient risk of CDI has been explored[12, 13, 14, 13], but population risk of CDI, to our knowledge, has not.

In this work, we build a patient mobility network for a large hospital using data spanning two years, and analyze the relationship between network centrality measures and CDI at the hospital unit level. Our approach takes into account the relationship of hospital-onset CDI – unit (population) susceptibility, mobility of patients, and environmental exposure. We describe a new practical clinical tool that can be used to quantify and visualize this relationship.

## 2. Methods

### 2.1 Human Subjects Protection

This proposal was reviewed and approved by the University of Rochester Human Subjects Review Board (protocol number RSRB00056930). Data were coded such that patients could not be identified directly in compliance with the Department of Health and Human Services Regulations for the Protection of Human Subjects (45 CFR 46.101(b)(4)).

### 2.2 Data Source and Study Population

De-identified patient electronic health record (EHR) data from a 739-bed hospital in New York State was acquired for a 2 year period from approximately January 2013 to December 2014 (n=209,694). These records include patient demographics, medication administration, ICD-9 diagnosis codes, lab results, and individual hospital unit admission data.

Admission for this study was defined as an admission of >24 hour duration to a hospital unit, either from the Emergency Department (ED) or directly. This definition was to ensure inclusion of patients who would be at risk of exposure to CDI as a hospital-onset case with a measurable outcome – i.e. a positive CDI lab result during their current admission. Multiple admissions for a single patient were included in this particular study as individual transfer and susceptibility data should still contribute to overall unit susceptibility, regardless of CDI case classification. Pediatric patients were excluded as they are not considered to be an at-risk population for CDI, and their inpatient units at this particular facility have little to no overlap with the adult inpatient units. Inclusion criteria of admission to a hospital unit beyond admission to the ED, admission of >24 hours, and patient age >18 years identified 72,636 eligible admissions in our 2-year data period.

A positive CDI outcome was determined via enzyme immunoassay (EIA) by nucleic acid amplification test (NAAT) and was further categorized as “hospital-onset” (positive lab occurred after a negative lab, or after patient had been admitted for 24 hours) or “community-onset” (positive lab occurred in the first 24 hours of a patient admission) as we did not have access to infection control data to further specify National Healthcare Safety Network (NHSN) standards for hospital-onset infection. Variables such as “length of stay”, and medication administration variables including antibiotic, proton pump inhibitor (PPI), and Histamine H2-receptor (H2) antagonists usage were adjusted to only capture data occurring prior to a patient’s first positive CDI test.

### 2.3 Network Construction

Relative inpatient unit admission date and time was calculated using EHR length-of-stay data within each patient admission, and was used to construct a 2-year patient mobility network. Weekly transfer rates from unit to unit were calculated using the following formula:

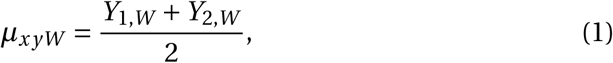

where *µ*_*x yW*_ = rate at which patients are be transfered between units *x* → *y* in week *W, x* = the transferring unit, *y* =the receiving unit, *W* = the calendar week, *Y*1,*W* = weekly data from year 1, and *Y*2,*W* = weekly data from year 2. Total weekly transfers from unit x to unit y for each calendar week (*W*) are calculated by adding the transfer sum of both years of weekly transfers, divided by two years of data.

Stochastic block modeling, a stochastic clustering method using a Bayesian approach, was used to determine non-degree correlated clusters of nodes in the network[16]. While this method can be useful for further analysis of community structure within a graph, this project uses it simply for visualization purposes and to examine units in groups based on their edge weights (transfer rates). Centrality measures, statistical values representing the connectedness of individual nodes in the network, were calculated for each network node (inpatient unit)[6, 7]. Network construction and centralities were performed using R Version 1.0.143, Python Igraph and GraphTool libraries, and Gephi Version 0.9.2.

### 2.4 Statistical Analysis

To capture patient risk in each inpatient unit, a logistic model was constructed using the training/test set method[17, 18], and an outcome variable of hospital-onset CDI as determined by our classification criteria. The data were randomly split 7:3, while ensuring our hospital-onset CDI case number was also randomly split by the same ratio. The resulting training set had *n*_*train*_ = 50, 845 patients with *c*_*train*_ = 321 CDI cases, and the test set had *n*_*test*_ = 21, 791 patients with *c*_*test*_ = 138 CDI cases.

Purposeful Model Selection [19] was used for the multivariate logistic model. This method was chosen in order to adequately capture our data and include variables of clinical significance that otherwise may have been excluded from the model in other selection methods. Daily mean individual patient risk (susceptibility) of CDI was calculated using the logistic model, and the overall mean inpatient unit susceptibility were calculated based on which patients were in each unit that day. Multivariate linear regression was conducted to determine association between overall mean unit susceptibility and unit centrality in the mobility network, with centralities being our predictor variables and unit susceptibility our outcome.

To capture inpatient mobility, unit susceptibility, and current infections in the hospital, a new and unique network centrality measure entitled “contagion centrality”, a term previously used to describe a cascading effect in interbank networks [20], was developed. Contagion centrality (CC), also known as the SFI Statistic (S = unit susceptibility, F = flow of patients, I = current infections) was calculated for each inpatient unit, every admission day:

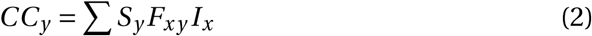

where *CC*_*y*_ = the contagion centrality of receiving unit, *S* = mean unit susceptibility, *F* =patient flow (transfer rate), *I* = current number of CDI infections, *x* = transferring unit, and *y* = receiving unit. The CC metric was evaluated through linear regression with an outcome of actual CDI cases. Regression, CC calculation, and other visualizations were performed using R Version 1.0.143 and Wolfram Mathematica Version 11.3.

## 3. Results

Our inpatient mobility network yielded 40 nodes (inpatient units) and 1,003 unique edges (inpatient unit-to-unit transfer pairs). The units with the highest mean incoming transfers are High Risk Post-Partum, High Risk Labor and Delivery, Cardiovascular Surgery, Trauma Surgery, and the Cardiac Intensive Care Unit (transfer rates of 7.84, 2.53, 1.66, 1.13, and 1.07 average patients/week, respectively). (Note: not all nodes are depicted in network graph figures and predictive tools – several inpatient units had little to no interaction with other units - meaning they did not have direct connections or transfers to other units in our hospital and did not contribute risk of CDI transmission to other units - deeming them unsuitable for our predictive tools.)

Our final multivariate logistic regression model yielded an AUC (Area Under the Curve) of 0.81, sensitivity of 0.75, and specificity of 0.71 at a threshold of 0.006 (0.6% of our patient sample of n=72,636 presented hospital-onset CDI; model variable selection can be viewed in Supplemental Table S1). All daily unit-specific susceptibilities were averaged over the full span of the dataset, yielding mean overall unit susceptibility for the 2 year period. Linear regression was used to compare the overall unit susceptibility to unit network centrality measures to determine whether unit susceptibility can be predicted by how connected a node is to other nodes in our mobility network via the transfer of patients (Table 1).

**Table 1:** **Mobility Network Centrality Multivariate Linear Regression**

| **Centrality Measure** | **Full-admission**<br>**variables<sup>a</sup>**<br>**(P Value)** | **Time-sensitive**<br>**variables<sup>b</sup>**<br>**(P Value)** |
| --- | --- | --- |
| In-Degree | 0.02 | NA<sup>c</sup> |
| Weighted In-Degree | 0.02 | NA |
| Closeness | <0.01 | <0.01 |
| PageRank | 0.02 | NA |

<sup>a</sup> Model does not contain time-sensitive medication administration variables – i.e. variables included if they occur at any point during a patient admission.
<sup>b</sup> Model contains time-sensitive medication administration variables occurring only prior to first positive CDI lab.
<sup>c</sup> NA values in time-sensitive model indicate the centrality measure was not statistically significant and left out of the model.

Table 1 presents two different susceptibility linear regression models and their associated centrality values. The regression model outcome for full-admission variables includes susceptibility variables from the logistic regression step prior to recalculation. This outcome captures the full patient admission as opposed to just the admission prior to the first positive CDI lab (time-sensitive variable outcome). Through Purposeful Model Selection, four centralities were associated with full-admission model susceptibility (In-Degree, Weighted In-Degree, Closeness, and PageRank), while the time-sensitive model susceptibility was only statistically significant with Closeness[6]. The difference between these two models lies in which part of the patient admission is most significant. Incoming transfers are best captured by Closeness, weighted in-degree, and in-degree, while the importance of both incoming and outgoing transfers is captured best by PageRank[6]. We conclude that PageRank centrality is useful for examining the full patient admission for all cause CDI cases. In contrast, the statistical significance of Closeness with our time-sensitive outcome suggests that the incoming edges (i.e. transfers into the patient unit) are important when analyzing susceptibility of hospital-onset CDI cases.

Generating the graph of our mobility network allows us to compare hospital unit centralities (Figure 1). Larger node size indicates a higher value of either Closeness or Susceptibility (depending on which mobility network is being viewed), while smaller node size indicates lower Closeness or Susceptibility. The coloring of the nodes indicates which communities (clusters) the inpatient units belong to as identified by stochastic block modeling[16] and is indicated by Groups A-G (group definitions can be seen in Supplemental Table S2). This method allows us to group units based on their connections to other nodes in the network, which can be useful when looking at transfer trends and patterns between groups. This method is simply used for visualization purposes in this study, and further clustering analysis would need to be conducted to determine community significance.

**Figure 1:**
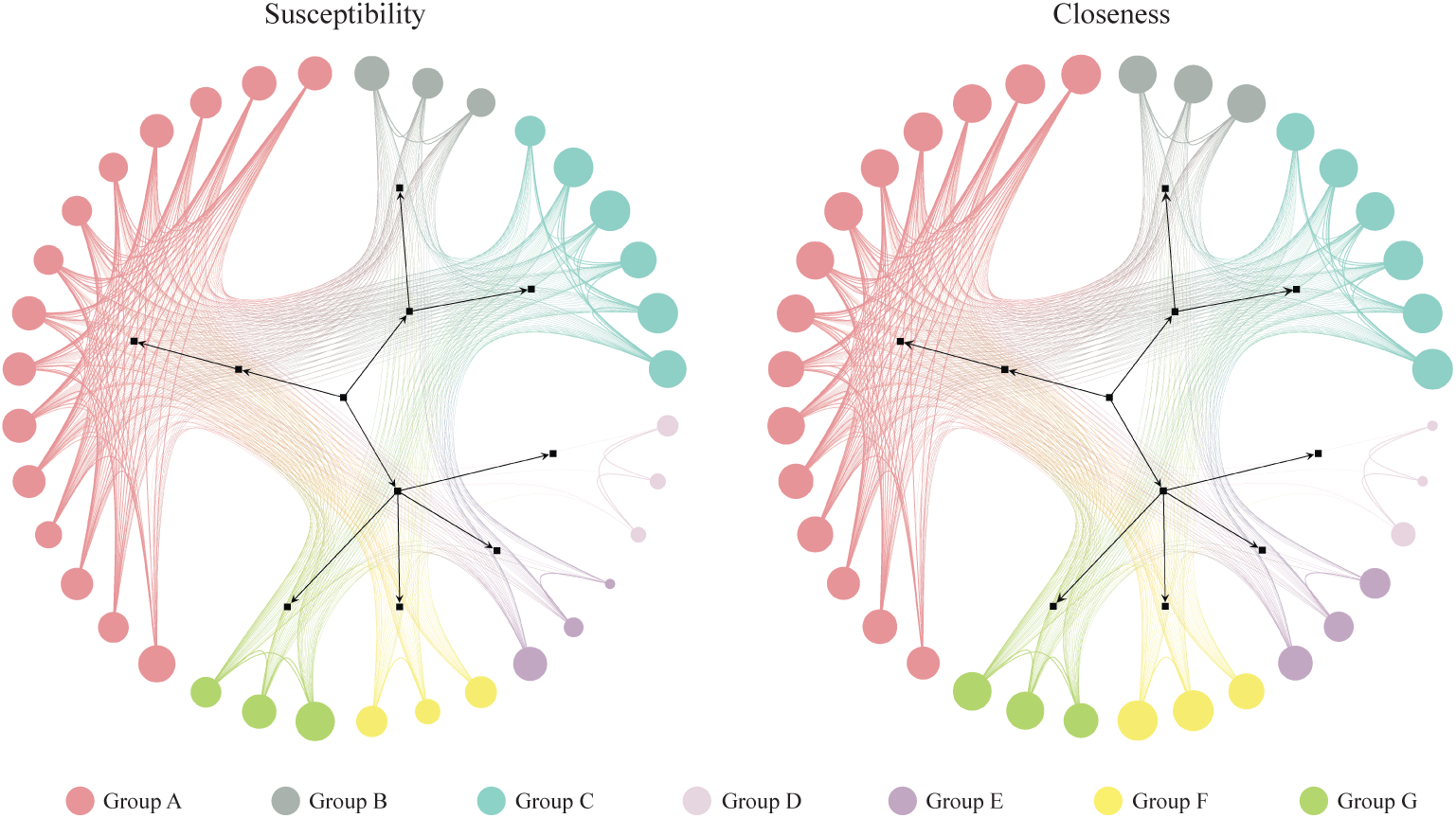
Patient Mobility Networks, Note: node size depicts normalized “Susceptibility” and “Closeness”, respectively.

Units with high Closeness tend to have higher CDI Susceptibility and vice versa, however clear exceptions to this relationship can be seen in Groups D, E, and F. Units in Group F may tend to be the last units in a patient’s journey (low Susceptibility), with high incoming rates and low outgoing transfer rates back to other units in our model (high Closeness). The units in Groups D and E may tend to transfer amongst each other without receiving or redistributing patients back into the network (low Closeness), but sometimes have patients with higher Susceptibility

To account for these exceptions while still capturing the important relationship of closeness centrality to risk in our other units, we developed the “Contagion Centrality” (CC) measure. The term “Contagion Centrality” was originally developed to explain the spread of financial instability in an interbank network[20] and is typically displayed in networks exhibiting a “spillover effect” or the rapid spread of fear of financial instability to predict bank failure. While a node with high financial CC is likely to influence other nodes it is connected to, our CDI contagion centrality is built simply from patient data within the current unit and from those units directly transferring into that unit. CC takes into account the incoming transfers captured by Closeness (F in our SFI Statistic), but adjusts for unit Susceptibility (S) and current infections (I). For example, a unit with high unit Susceptibility (S) who receives many patients from units who currently have CDI present (high F and I) would in turn have a high CC and high risk of CDI appearing in their unit, as well. The units with highest overall CC (normalized between 0-100) are the acute medicine units 4, 1, 3, and 2 (CC = 100, 99.9, 99.9, and 99.7, respectively).

CC metric validation through linear regression with the outcome of actual CDI cases yielded statistical significance (P value <0.05), thus confirming our hypothesis that inpatient transfers are associated with hospital-onset CDI. Difference plots comparing daily CC and actual CDI cases determined the appropriate cut-off points for each unit for which day prior to CDI our model is most predictive. The mean predictive period was 3.33 days with the majority of units falling between 1-6 days prior to the CDI case.

Highlighting the importance of incoming inpatient transfers for unit-wide susceptibility of hospital-onset CDI provides infection prevention teams and clinicians with more information, but lacks the precision needed to actually prevent these infections on a unit-by-unit basis. Plotting deconstructed CC not only provides hospitals with a daily CC value by unit, but informs them of factors contributing to that value – i.e. flow of infection or unit-wide susceptibility (Figure 2).

**Figure 2:**
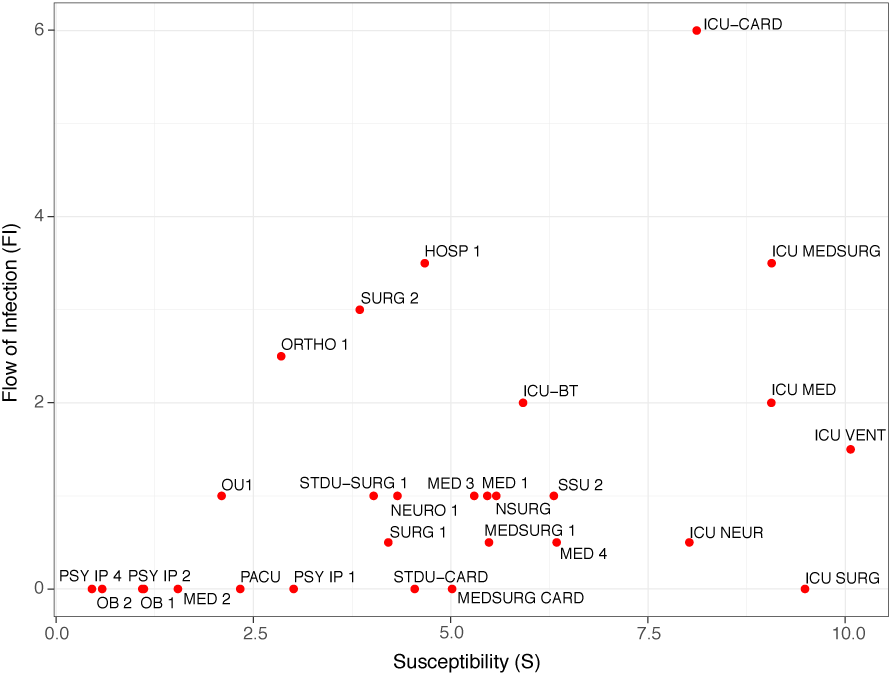
Daily Contagion centrality calculation plot example.

Viewing the information in this way allows clinicians and infection preventionists to not only track all units at one time, but to also see which factors are contributing to an increase or decrease in CC. Units with higher FI (y-axis) may have an increased CC due to more transfers or potential transfer of infection into that unit. Units with higher S (x-axis) may have an increased CC due to the patients they currently have in their unit that day and their susceptibility to infection.

Supplementing the calculation plot is an additional active surveillance tool showing an overall plot of unit change (units with the most positive or negative changes are highlighted and labeled), and subplot of individual unit weekly change (Figure 3).

**Figure 3:**
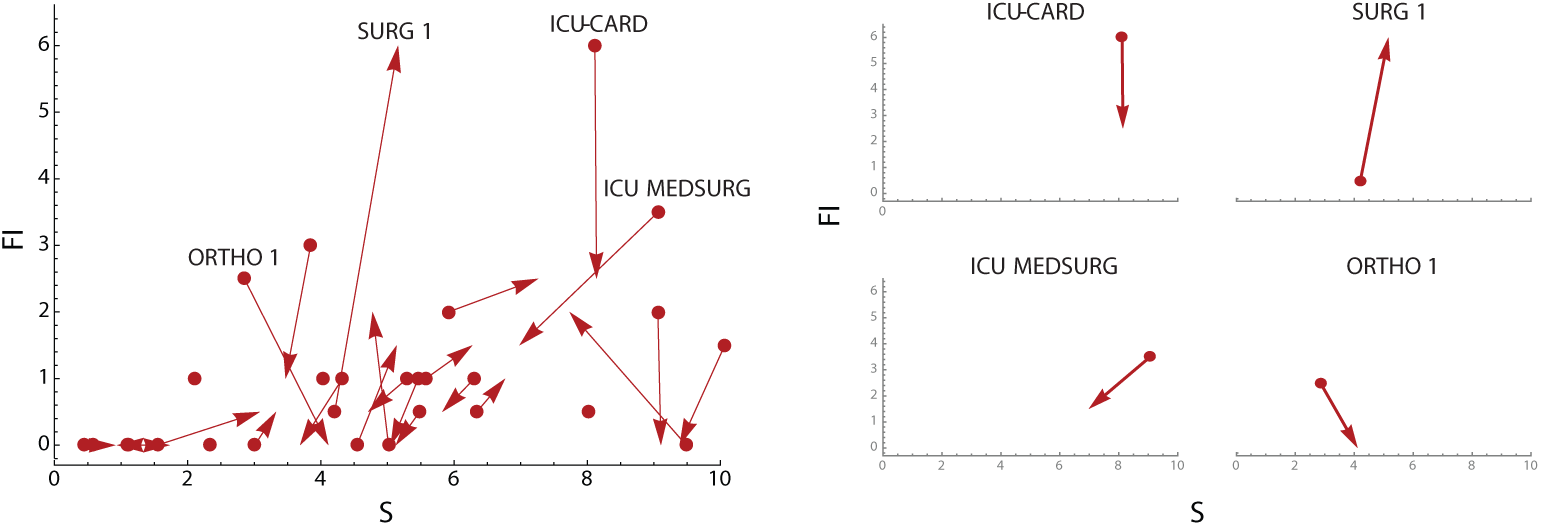
Weekly Contagion Centrality change plots, Note: overall plot (left) shows change in all inpatient units in a 7-day period, and unit plots (right) show change in select individual inpatient units in a 7-day period. S, susceptibility; FI, flow of infection.

Plotting change allows infection preventions and outcomes specialists to observe weekly change in all units at one time, as well as isolate each unit and look at increases or decreases in CC on an individual basis, all while still being able to see contributing factors to a high or low CC. Use of these tools together provides unique and active surveillance to prevention teams, and can help hospitals prepare for situations of heightened CDI infection risk.

## 4. Discussion

To our knowledge, this is the first multifactorial clinical tool for the prediction and surveillance of hospital-onset CDI. A strength of our study is the inpatient mobility network construction and analysis. As previously mentioned, literature tends to focus primarily on inter-hospital transfers as opposed to intrahospital transfers, and we have shown that incoming inpatient transfers are statistically associated with hospital-onset CDI. An additional strength of this tool is that it can be tailored to fit different facilities, and possibly even different infections. It is designed to identify units with shared opportunities of contact, so for now, it is limited to organisms transmitted via contact.

Limitations of our study include limited medication administration data stemming from a single facility. Additional categorical variables were originally created to indicate if a patient had received antibiotics in the 14- and 30-day time periods prior to their current admission, but was not ultimately used in this study. It is likely patients could have received antibiotic, H2 antagonist, or PPI medication from outside providers in the weeks leading up to their inpatient admission, posing susceptibility not captured by our data. Another limitation is the lack of formal infection prevention lab classification. Hospital-onset CDI data was unavailable for patient-specific admissions, thus the general classification of hospital-onset cases simply being those occurring after 24 hours of admission or after a previously negative lab result, and may result in a slightly higher number of reported hospital-onset CDI cases for this facility. Our last limitation is situational – if a unit’s susceptibility is low, but CDI rates in that unit are still high, it is possible an outside factor such as decreased hand hygiene or environmental contamination could be contributing to hospital-onset CDI cases. Further analysis and surveillance would be required to pinpoint the cause(s) contributing to the higher rate.

Future studies may benefit from network analysis of unit community clustering to determine the risk of hospital-onset CDI on communities of units instead of single units, as well as further calibration analysis to determine probability of prediction by unit. Implementation and trial of Contagion Centrality and the associated visualization tools in a hospital or facility would further validate our model and highlight facility-specific changes that should be made to better capture CDI outbreak risk and allow for intervention development to measure and reduce unit CC and risk of hospital-onset CDI. Each hospital unit in our data tends to have different trends and patterns of CC when examining increases leading up to actual CDI cases. Our recommendation is that while hospital trial is needed to pinpoint unit-specific periods of highest risk of CDI following an increase in CC, increasing infection prevention methods and awareness for the following 7 days is a good place to start.

It is important to recognize the knowledge gap between inpatient mobility and risk of infection, as well as the need for more prospective infection prevention tools. This study shows the significant association between inpatient transfer and unit risk of CDI, and provides the first attempt to our knowledge to actively measure and predict this occurrence.

## Acknowledgments

This work was partially funded by the University of Rochester Clinical and Translational Science Institute, supported in part by grants UL1 TR002001, and TL1 TR002000 from the National Center for Advancing Translational Sciences (NCATS), a component of the National Institutes of Health (NIH). Additionally, this work was funded by the Translational Biomedical Science PhD Program at the University of Rochester School of Medicine and Dentistry through support from grants 1014095 BWF from the Burroughs Wellcome Fund Institutional Program Unifying Population and Laboratory Based Sciences and TL1 TR000096 Training Grant from the National Center for Advancing Translational Sciences of the National Institutes of Health. The content is solely the responsibility of the authors and does not necessarily represent the official views of the National Institutes of Health (NIH) or Burroughs Wellcome Fund (BWF).

We would also like to thank Dongmei Li and Solomon Abiola at University of Rochester Medical Center for discussions regarding the statistical analyses.

